# Concurrent selection of internal goals and external sensations during visual search

**DOI:** 10.1101/2025.02.25.640053

**Authors:** Baiwei Liu, Freek van Ede

**Author notes:** Correspondence: Baiwei Liu, Freek van Ede.

## Abstract

Flexible goal-directed behaviour relies on the selective processing of internal goal representations and external sensations. Yet, internal and external selection processes have classically been studied in isolation, leaving us in the dark how internal and external selection processes are coordinated in time to support behaviour. To address this, we developed a novel visual-search task in which we could simultaneously track selection among internal search goals held in working memory and external search targets in the environment. Capitalising on sensitive gaze and neural markers of internal and external visual selection, we provide proof-of-principle evidence in humans that internal and external selection processes do not necessarily take turns in a strictly serial manner, but can develop concurrently. We further show how concurrent internal and external selection processes are associated with largely non-overlapping neural activity patterns in the human brain, and how these processes can be performed effectively even when engaging opposite spatial locations in working memory and perception. These findings challenge views portraying brain states as being either internally or externally focused and bring new insight into how internal and external selection processes work together to yield efficient search behaviour.

## INTRODUCTION

Goal-directed behaviour fundamentally relies on the selection of internal goal representations and goal-relevant external sensations, such as during search where the activation of specific search goals guides the selective processing of incoming sensations. While selection processes have been studied extensively for selection among the external contents of perception ^1–5^ or among the internal contents of working memory ^6–9^, there has remained a relatively scarcity of studies that considered how internal and external selection processes work together ^10–12^ – despite behaviour often relying on both. This has left us in the dark regarding the foundational question how internal and external selection processes are coordinated in time to support efficient goal-directed behaviour.

One possibility is that selective internal and external visual processes necessarily ‘take turns’, with our attentional focus moving serially between sampling internal goal representations (objects we search for) and sampling external information (objects we search among). Such a scenario fits with views in which, at any given moment, the brain is either focused *externally* toward the content of perception and action or *internally* toward our memories and thoughts ^13–17^. In such a framework, when we rely on the joint selective processing of both internal and external information to guide our behaviour, we must switch focus between them in a serial manner ^18–23^. Alternatively, given the prevalence of everyday tasks that jointly rely on the selective processing of external and internal information, it is conceivable that the brain may have developed ways to direct its attentional focus to relevant internal and external information concurrently – and in a way that minimises interference between these processes.

Here we studied the temporal coordination of internal and external selection dynamics in the context of visual search. Search is a ubiquitous daily task that exemplifies our reliance on joint internal and external selective processing, as finding what one is looking for relies on the selective activation of *internal* search goals to guide the selective processing of *external* incoming sensations. Yet, while it is well appreciated that search occurs at the confluence of selective internal and external visual processes ^5,22,24–27^, how internal and external selection processes are coordinated during search has remained elusive. This is owing, at least in part, to methodological challenges of isolating and tracking internal selection processes while concurrently processing external visual inputs.

We developed a search task that enabled us to independently and simultaneously track visual focusing among internal search goals held in working memory and the external objects of search. Capitalising on sensitive gaze and neural markers of internal and external visual selection, we provide proof-of-principle evidence in humans that internal and external selection processes need not take turns in a strictly serial manner, but can proceed concurrently. We further show how concurrent internal and external selection processes are associated with largely non-overlapping neural activity patterns in the human brain and can be deployed effectively even when engaging opposite spatial locations in working memory and perception.

## RESULTS

Human volunteers performed a search task that jointly required the internal selection of a search goal held in working memory and external selection of the search target in the external search display (**Fig. 1a**). Participants held two coloured objects in working memory until a cue (a colour-change of the central fixation cross) prompted participants to selectively search for the colour-matching memory object among four greyscale objects (the search display). The task was to indicate the location of the search target in the search display using a key press. The search display became available for inspection together with the memory cue, inviting immediate utilisation of the selected internal search goal. Because the search display always contained both memory objects (together with two novel objects), the selection of the cued internal search goal was a prerequisite for external search. This enabled us to address whether internal and external selection processes necessarily take turns (completing selection of the internal search goal *before* externally searching it) or can proceed concurrently (starting to guide external search from the moment the internal process of search-goal selection is initiated).

**Figure 1.**
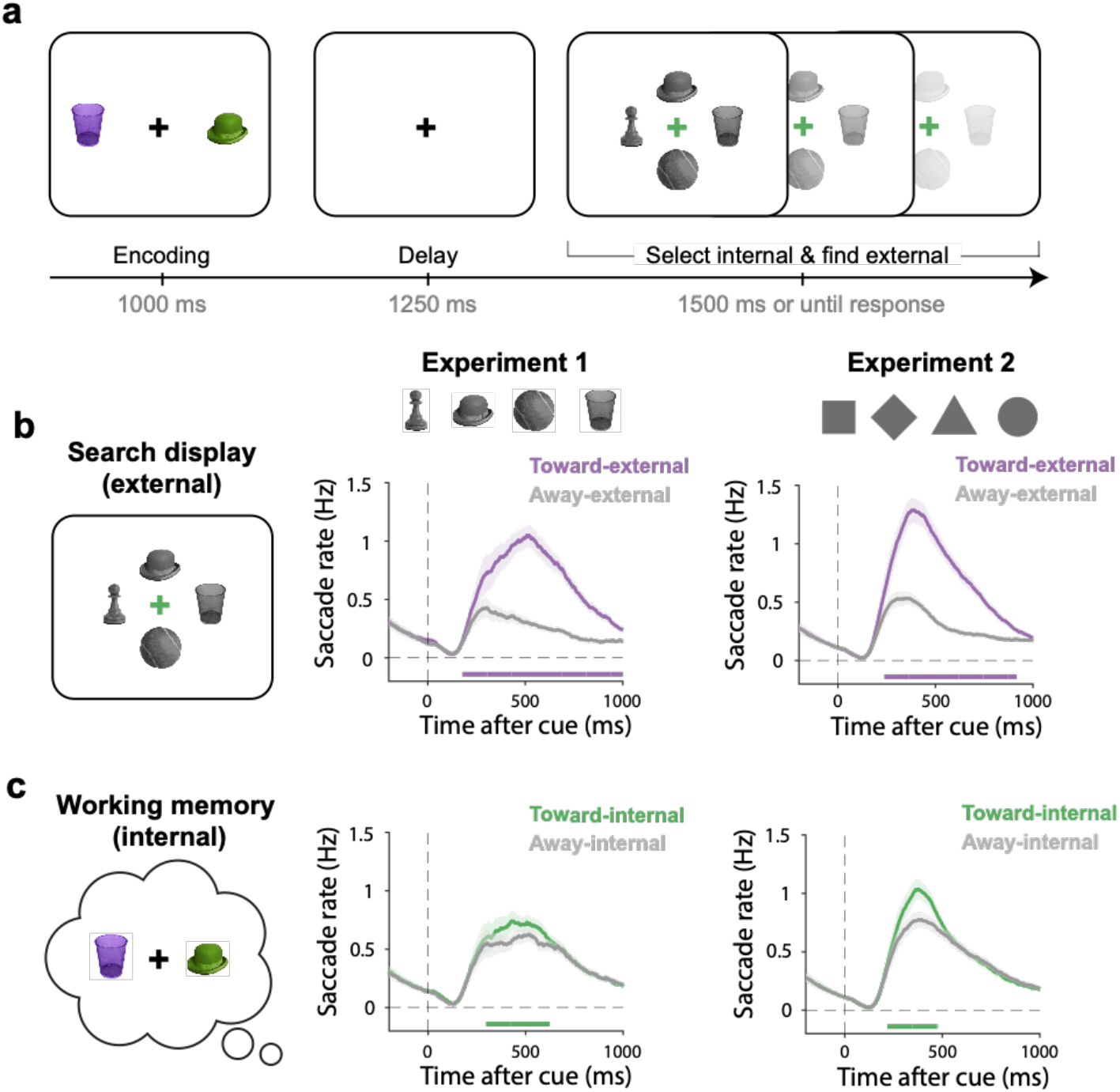
Eye-movements track the selection of internal search goals and external search targets. **a)** Schematic of the memory-guided visual-search task. Participants encoded two potential search goals into working memory until a colour cue indicated the memory object whose match had to be located among four grey-scale objects in the external search display. Internal and external locations associated with the cued object were manipulated independently. Experiment 1 used images of real-world objects while Experiment 2 used simple shapes (stimuli examples shown in panel b, see also **Supplementary Fig. 1). b)** Time course of external selection as measured by saccade rates toward and away from the location of the search target in the external search display. c) Time course of internal selection, measured by saccade rates toward and away from the memorised location of the cued search-goal object held in working memory. All time courses show mean values, with shading indicating ±1 SEM calculated across participants (n=25 in Experiment 1; n=50 in Experiment 2). The thick horizontal lines in the time course plots indicate the significant temporal clusters of the difference in saccade rates toward vs. away from the internal and external location of the cued memory object (cluster-based permutation P < 0.05).

We ran two experiments. After validating our task and eye-movement findings with images of real-world objects in Experiment 1, we sought to replicate and extend our findings in Experiment 2. In Experiment 2 we boosted sensitivity by using simpler shape stimuli, collecting more trials per participant, doubling the number of participants, and including EEG measurements. In both versions of the task, participants performed well above the chance, with an accuracy of 87±2% (mean±SE) in Experiment 1 and 83±1% in Experiment 2. Participants also responded well within the maximum response time of 1500 ms with an average reaction time of 871 ±18 ms in Experiment 1 and 785±12 ms in Experiment 2.

### Eye-movements track the selection of internal search goals and external search targets

In our task, we independently manipulated the location in working memory of the cued memory object and the location on the screen of the external search target. This enabled us to independently track the selection of internal search goals and external search targets. We first turned to spatial biases in eye-movements, which have recently been demonstrated to track not only external, but also internal selection within the spatial layout of visual working memory ^28–31^. Before turning to the temporal coordination of internal and external selection processes, we first verify the utility of eye movements for tracking both internal (search-goal) and external (search-target) selection, in our task that relies on both.

Not surprisingly, *external* selection of the target object in the external search display could be robustly tracked through the direction of eye movements. Specifically, we observed a higher rate of saccades in the direction of the search-target in the external search display as opposed to in the opposite direction (**Fig. 1b**). This was the case both in Experiment 1 (cluster P < 0.001) and Experiment 2 (cluster P < 0.001).

Critically, eye movements also tracked *internal* selection of the cued search goal held within the spatial-layout of visual working memory (whose memorised internal location was manipulated independently from the external location of this object in the search display). This was again the case in both experiments (**Fig. 1c**, cluster Ps < 0.001). These findings build on recent demonstrations that the selection of visual objects held within the spatial layout of working memory can be read-out from spatial biases in eye movements ^28–31^. We now show that this is the case *even* in the presence of an external search display. Having established this, we next turn to our central question regarding how these internal and external selection processes are coordinated in time.

### Internal and external selection unfold concurrently during search

Figure 2. zooms in on the internal and external selection markers of interest, by showing the difference between toward and away saccades, as defined in reference to the memorised internal location of the cued memory object (internal) or its external location on the screen (external). Consistent with ample prior work ^7,29,32–36^, we show that it takes approximately 200 ms to deploy internal selective attention after the memory cue, as here signalled through spatial biases in eye movements to the cued memory object (**Fig. 2a**, green lines). Critically, our data show that it does not take *another* 200 ms to subsequently initiate selection of the external memory target on the screen (**Fig. 2a**, purple lines). Instead, selection of the external search target strongly overlaps in time with selection of the internal search goal held in working memory.

To compare the observed temporal profiles associated with internal and external selection, we normalised the spatial saccade modulations as a percentage of their maximum value (**Fig. 2b**), exposing their striking concurrence in time. To quantify this temporal concurrence, we deployed a jackknife-based analysis ^37^ on the latency of our spatial saccade markers of internal and external selection (quantified as the first value reaching 50% of the peak). We found no significant differences in latency in neither experiment (ps > 0.6). Moreover, a Bayesian analysis showed moderate evidence in favour of the null hypothesis of no latency difference between the internal and external selection time courses, in both Experiment 1 (BF01 = 4.3) and Experiment 2 (BF01 = 5.8).

**Figure 2.**
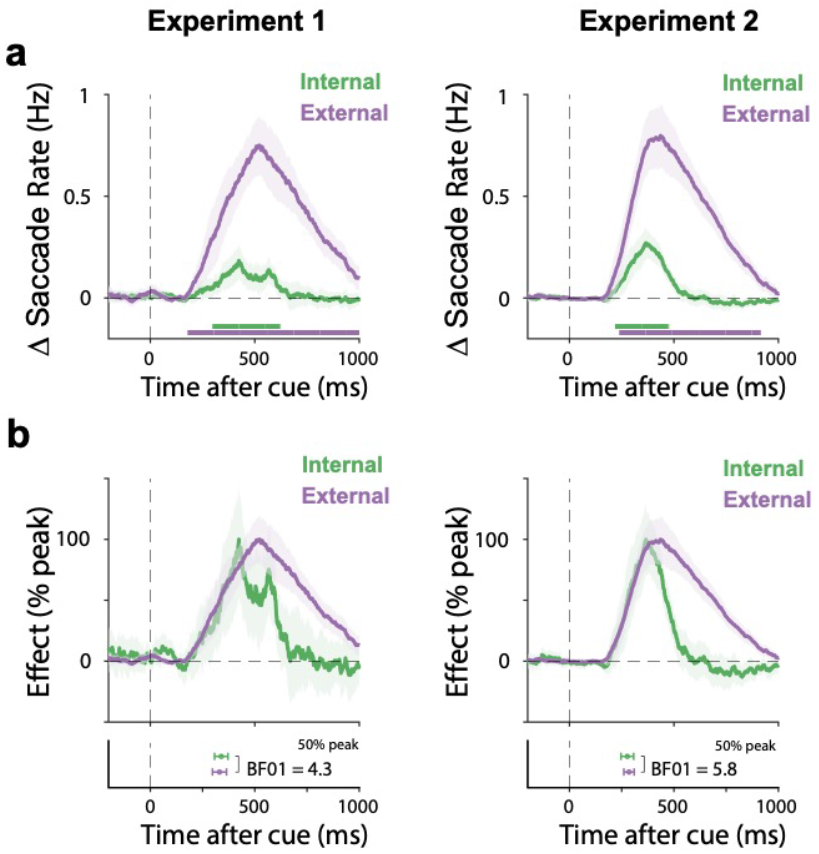
Internal and external selection unfold concurrently during search. **a)** Time courses of spatial saccade modulations associated with internal and external selection in Experiment 1 (left) and Experiment 2 (right), as operationalised as saccade rates toward vs. away from the respective locations of the cued search target in working memory (internal) and in the search display (external). b) The same data as in panel a, following normalisation as a percentage of the maximum value. Bottom panels show onset latencies, quantified as the first sample reaching 50% of the peak. Time courses show mean values, with shading indicating ±1 SEM calculated across participants (n=25 in Experiment 1; n=50 in Experiment 2). The thick horizontal lines in the time course plots indicate the significant temporal cluster (cluster-based permutation P < 0.05). Error bars on the onset latencies in the bottom panels were estimated using a Jackknife approach and show mean ± the 95% confidence interval. Bayes factors (BF01) indicate evidence in favour of the null hypothesis of no difference.

The concurrence of internal and external selection held even when only considering trials in which internal and external selection occurred in perpendicular axes and in trials in which the two memory objects competed along the same axis in the search display, ruling out low-level explanations for the observed concurrence of internal and external selection in our task (see **Supplementary Results** and **Supplementary Fig. 2**).

### Internal and external selection rely on largely non-overlapping neural activity patterns in the human brain

In Experiment 2, we supplemented our eye-movement measurements with EEG measurements. This enabled us to additionally address whether the observed concurrent selection of internal and external visual targets is mediated by shared or distinct neural activity patterns in the human brain.

Again, we utilised our independent manipulation of internal and external object locations in working memory and in the search display. This enabled us to independently train classifiers to decode information regarding the internal or external locations associated with the cued object from multivariate patterns of EEG activity. To do so, we employed a time-resolved multivariate classification to broadband EEG data. Akin to the eye-movement data, we could robustly decode internal (search-goal, cluster P = 0.002) and external (search-target, cluster P < 0.001) locations associated with the cued object, as shown in **Figure 3a**. Normalizing these signals (**Fig. 3b**) again showed considerable overlap in time.

**Figure 3.**
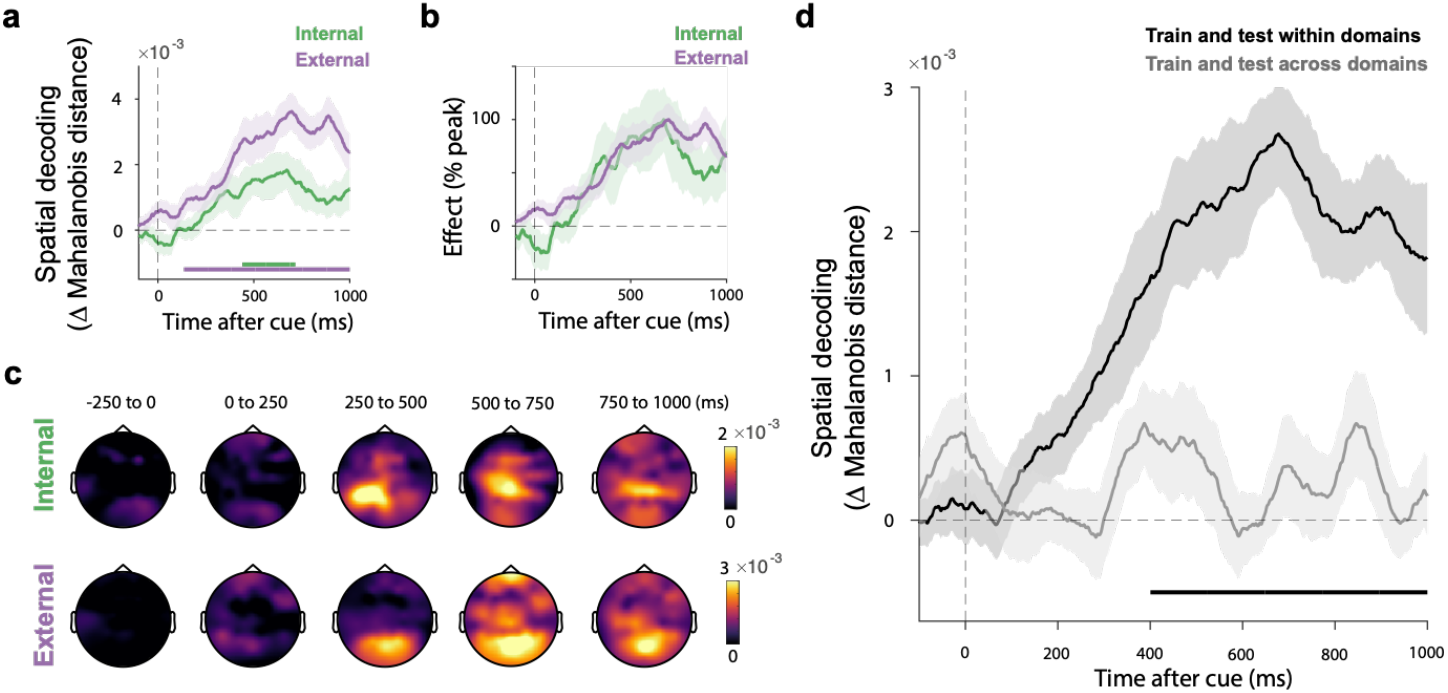
Internal and external selection rely on largely non-overlapping neural activity patterns in the human brain. **a)** Time courses of multivariate decoding of the spatial direction associated with internal and external selection, expressed as the difference in Mahalanobis distances (non-matching minus matching, yielding positive decoding values). Thick horizontal lines indicate significant temporal clusters for each condition (compared to zero, cluster-based permutation P < 0.05). **b)** Time courses of decoding normalised as a percentage of their maximum value. **c)** Topographies associated with the spatial decoding of internal and external selection as a function of time. Decoding topographies were constructed using an iterative searchlight approach. **d)** Time courses of within-domain decoding (e.g., training and testing classifiers separately within internal and external selection) and across-domain decoding (e.g., training on internal selection and testing on external selection, or vice versa). All time courses show mean values, with shading indicating ±1 SEM calculated across participants. The thick horizontal lines in the time course plots indicate significant temporal clusters (cluster-based permutation P < 0.05). In panel d, the significant temporal cluster indicates the difference between within- and across-domain decoding (cluster-based permutation, p < 0.05).

To address whether these co-developing selection processes arise from shared or distinct neural activity patterns, we performed two additional analyses. First, we conducted a searchlight analysis (as employed in ^11,35,38,39^), iteratively running the decoding analysis on subsets of EEG electrodes to identify scalp regions contributing to internal and external spatial selection. This provided tentative evidence that internal and external selection processes, while unfolding concurrently, have distinct neural origins. Patterns associated with internal selection appeared more anterior than those associated with external selection (**Fig. 3c**).

To more directly quantify the shared vs. distinct nature of the neural activity patterns associated with internal and external selection, we next performed a cross-domain decoding analysis. We trained a classifier to distinguish object direction in the internal or external space, and tested the classifier within the same domain (train-internal->test-internal, train-external->test-external) or across domains (train-internal->test-external, train-external->test-internal). As shown in **Figure 3d**, while we observed robust decoding when training and testing within the same domain, we found a striking lack of generalisation across internal and external domains (no significant cluster found: *P*s > 0.2). This was reinforced by significantly better decoding within vs. across domains (cluster *P* < 0.001). These data provide direct evidence for the use of distinct, largely non-overlapping neural activity patterns associated with the co-occurring internal and external selection processes in our task.

### Internal and external selection can be deployed in opposite directions without a cost

We finally looked for potential interactions – or lack of such interactions – when internal and external selection occurred in the same or opposite direction, relative to perpendicular directions that here served as a baseline. Strikingly, as shown in **Figure 4**, when internal and external selection occurred in opposite directions (e.g., internal left, external right), performance was not worse than the baseline condition in which internal and external selection occurred in perpendicular axes (e.g., internal left and external top). This was the case neither for accuracy (Experiment 1: t(24) = 0.53, p = 0.6, d = 0.11, BF01 = 4.18; Experiment 2: t(49) = 0.58, p =0.57, d = 0.08, BF01 = 5.54), nor for reaction time (Experiment 1: t(24) = 1.2, p = 0.24, d = 0.24, BF01 = 2.48; Experiment 2: t(49) = −1.09, p = 0.28, d = −0.15, BF01 = 3.73). Thus, search appeared equally effective no matter whether internal and external selection occurred in perpendicular (un-correlated) or opposite (anti-correlated) directions. Adding to our EEG results that showed largely non-overlapping activity patterns governing internal and external selection, these behavioural data suggest how internal and external selection can be deployed opposite directions, without necessarily incurring a cost.

**Figure 4.**
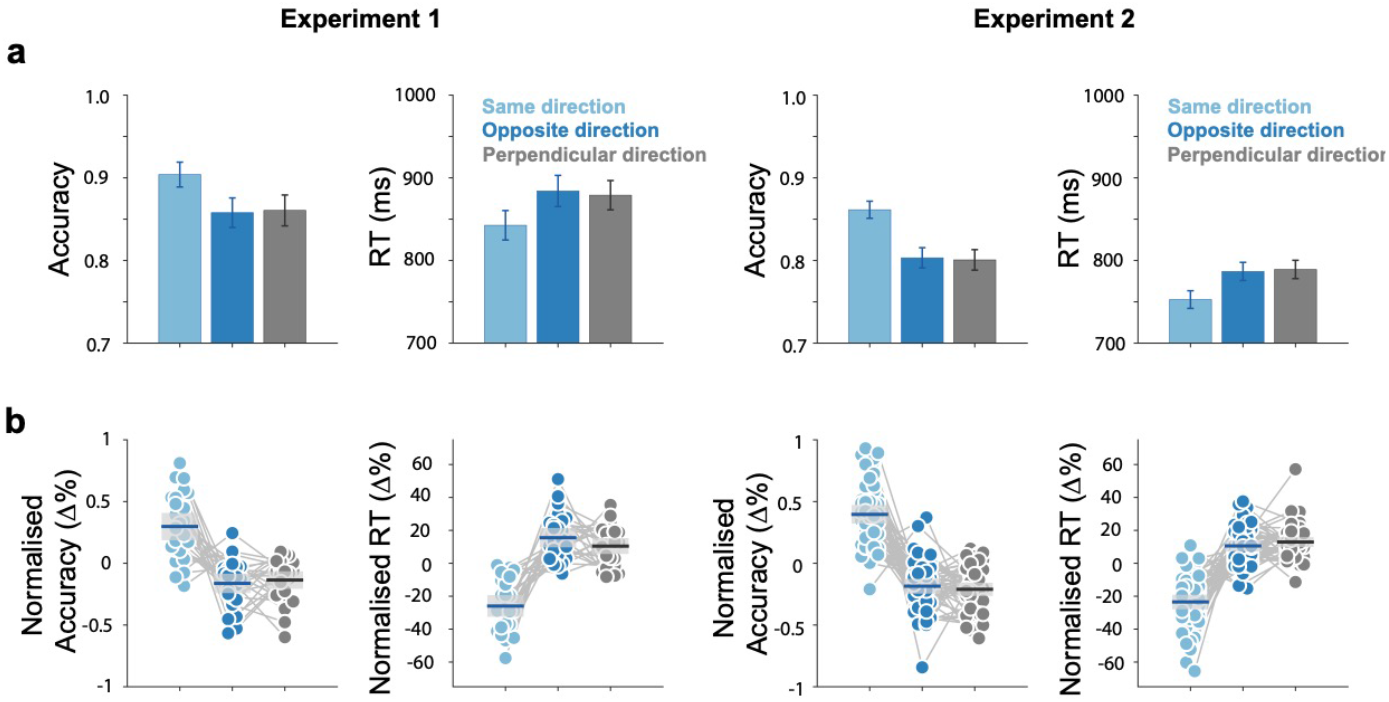
Internal and external selection can be deployed in opposite directions without a cost. **a)** Trials were categorized into three conditions based on whether internal and external selection occurred in the same, opposite, or perpendicular directions relative to the central fixation cross. Accuracy and reaction time of the three conditions in Experiment 1 (left) and Experiment 2 (right). Data depict mean performance with error bars indicating ±1 SEM. b) Normalized performance (percent change from the mean) with individual data points overlaid and shadings representing 95% confidence intervals.

At the same time, our behavioural data showed how internal and external selection-through-location were not fully independent: when internal and external selection occurred in the same direction (e.g., internal left and external left), search was selectively facilitated. This was reflected in higher accuracy (Experiment 1: t(24) = 5.24, p < 0.001, d = 1.05; Experiment 2: t(49) = 10.18, p < 0.001, d = 1.44) and faster reaction times (Experiment 1: t(24) = −7.45, p < 0.001, d = −1.5; Experiment 2: t(49) = −10.49, p < 0.001, d = −1.48) compared to the same baseline condition in which the selected memory object and the external search target in the display were on perpendicular axes. Thus, even if there is no cost associated with deploying internal and external selection in opposite (as opposed to perpendicular) directions, there is nevertheless a benefit when internal and external selection match in direction.

## DISCUSSION

Our data bring new insight into how internal and external selection processes combine to yield efficient goal-directed behaviour, as we studied in the context of visual search. We provide proof-of-principle evidence in humans that internal and external selection processes do not necessarily take turns in a strictly serial manner but can develop concurrently. We further show how concurrent internal and external selection processes rely on largely non-overlapping neural activity patterns in the human brain and how internal and external selection can be deployed effectively even when engaging opposite spatial locations in working memory and perception. These findings challenge views portraying brain states as being either internally or externally focused, and reveal new insights into how internal and external selection processes combine to yield efficient search behaviour.

A large body of prior research has delineated the mechanisms and principles governing selective information processing (selective attention) as deployed either among the content of perception ^1–5,26^ or among the content of working memory ^6–9^. Additional studies have directly compared the two ^6,7,32,40^. Yet, it has remained surprisingly underexplored how such selective processes are coordinated in tasks that rely on both internal and external selection processes. Complementing prior studies that investigated the mechanisms supporting switching of attention between contents in perception and working memory ^10–12,19,21,41^, we uniquely studied whether internal and external selection can be deployed jointly, or necessarily take turns. Contrary to the intuition that the selection of an internal search goal necessarily precedes the selection of the search target in the external world, in our task, the process of selecting the relevant search goal and using it to guide the selection of the relevant external target co-developed. While we do not claim that internal and external selection will *always* concur, our data provide proof-of-principle that they *can* concur and are thus not inherently serial whereby internal selection has to be completed before external selection can start.

In our task, internal selection was a prerequisite for guiding external selection. The search display always contained both memory objects and participants were required to selectively find only the object that was cued through colour (with object colour exclusively existing in memory). This aspect of our task makes our findings all the more remarkable. Indeed, given that our task required internal selection to guide external selection, why did we not observe any delay between our internal and external selection markers? We posit that such a delay would only be expected when assuming that the internal search-goal selection – a process that consumes time – has to be *completed* before it can guide external search. Instead, our findings suggest that the process of internal selection starts guiding external search (cf.^5,25,42–46^) as soon as the process of internal search-goal selection is initiated (even if itself still developing), leading internal and external selection to unfold together.

Besides their concurrence in time, our data also show how internal and external selection processes rely on complementary neural activity patterns in the human brain, with internal visual selection appearing more anterior than external visual selection processes (consistent with ^32,47,48^). Furthermore, multivariate pattern analyses revealed clearly decodable selection patterns in both perception and working memory (see also ^11,49^), but with little cross-domain generalisation between internal and external spatial selection (see also ^7^). This suggests distinct neural coding for internal and external selection, akin to the type of orthogonal neuronal coding that has been reported between working-memory contents and external distractors ^50^, for working-memory contents before and after their selection ^7^, or for different streams of information held in working memory ^51^. Critically, such largely non-overlapping neural coding may be a prerequisite for the two processes to run concurrently without interfering with each other (see also ^52^). Indeed, if internal and external selection engaged the same overlapping neural activity patterns, their concurrent deployment would be challenging.

Our behavioural data corroborated the relative independence by which internal and external selection can be deployed: showing how internal and external selection could be employed effectively even when directed in opposite (compared to perpendicular) directions in working memory and perception. At the same time, we observed a selective benefit when internal and external selection aligned in perception and working memory. One possible explanation for this finding is that the spatial gaze bias associated with internal selection also brought the relevant external target closer to the fovea in these trials, in turn facilitating search.

We leveraged spatial gaze biases and multivariate EEG patterns track internal search-goal and external search-target selection. These markers each bring excellent time resolution and converged on concurrent selection patterns. In both markers, the internal selection process started approximately 200 ms after cue onset, consistent with ample prior research from complementary tasks ^7,29,32–36^. Critically, both markers also clearly showed that it does *not* take another 200 ms to then deploy selective attention to the external memory target on the screen – as may be expected from a strictly serial deployment of internal and external attention. Though it is notoriously challenging to prove zero delay, our findings are unambiguous in showing how internal and external selection processes – as reflected in our markers – emerge early after the cue and co-develop in time. Even so, we cannot rule out that our markers failed to capture additional components of internal and external selection processes that operate in a more serial manner, or that internal and external selection processes engage in more rapid turn-taking, within the timescale at which we show their co-development. Future work using invasive neural recordings across visual, frontal, and subcortical areas are required to assess these possibilities.

By experimentally orthogonalizing internal and external selection processes, our study brings a novel approach for delineating how internal and external processes work together to yield efficient search behaviour. This approach hold promise for extending it to other human behaviours that, like search, rely on the selective processing of relevant internal goal representations and external sensations.

## METHODS

We conducted two experiments to examine the temporal dynamics of internal and external selection processes during memory-guided visual search. In Experiment 2, we replicated and extended the findings of Experiment 1 by simplifying the stimuli, doubling the number of participants, adding more trials, and incorporating EEG recordings.

### Ethics

Experimental procedures were reviewed and approved by the Ethics Committee at the Vrije Universiteit Amsterdam. Participants provided written informed consent before the experiment and were compensated €10 per hour (or the equivalent in credits) for their time.

### Participants

In Experiment 1, 25 participants (18–33 years; 9 male, 15 female, 1 non-binary; 24 right-handed; 5 with corrected-to-normal vision) completed the task. Experiment 2 was conducted in two parts. For the first part, 25 participants (18–28 years; 4 male, 21 female; 25 right-handed; 9 with corrected-to-normal vision) were recruited to perform the eye-tracking task alone. For the second part, an additional 25 participants (18–30 years; 6 male, 39 female; 23 right-handed; 7 with corrected-to-normal vision) completed the same task with both eye-tracking and EEG recordings. The EEG data from one participant had to be excluded due to corrupted EEG files that could not be read-in during analysis. The basic sample size of 25 was set a-priori based on prior studies from our lab, which successfully tracked spatial selection in working memory using similar eye-tracking and EEG outcome measures. Because the two parts of Experiment 2 differed only in whether or not EEG was also collected, we pooled their behavioural and eye-tracking data to maximise sensitivity.

### Experimental design

In both experiments, participants performed a memory-guided visual search task (**Fig. 1a**). Each trial began with the presentation of two objects (potential search goals) of a different colour. These objects appeared left and right or top and bottom of a central fixation cross (at 6 degrees in Experiment 1 or 5 degrees in Experiment 2). Participants were instructed to encode these objects into working memory and to wait for a colour cue that instructed them to selectively search one of them in a visual search display.

In Experiment 1, the encoding display remained visible for 1000 ms. In Experiment 2, which used simpler shapes, the encoding time was reduced to 250 ms to fit more trials into the same duration. In both experiments, after a 1250 ms retention interval, the search display appeared simultaneously with a change in the colour of the fixation cross. This colour-change of the central fixation cross served as the memory cue prompting participants to select the colour-matching object from working memory as their search goal. Participants were instructed to find this memory object in the search display, from which all colour was removed. The search display always contained both objects that had been held in working memory plus another two filler objects. Objects in the search display were always displayed in greyscale, such that the use of the colour cue had to engage working memory.

The four objects in the search display were arranged at the top, bottom, left, and right around the central fixation cross (at a radius of 2 visual degrees in Experiment 1 or 1.5 visual degrees in Experiment 2). Participants responded by pressing one of four arrow keys corresponding to the target’s location in the external search display. Participants were required to indicate the location of the *cued* memory object and to ignore the other, uncued memory object in the search display. To encourage rapid responding, the search display gradually faded and disappeared entirely after 1500 ms, which also served as the maximum response time. After the response, the stimulus selected by the participant would turn white for 250 ms together with feedback (“0” for wrong, or “1” for correct) printed above the fixation cross. The next trial would start after an inter-trial interval randomly drawn between 500 and 1000 ms.

Critically, the location of the cued memory object during memory encoding and the location of this object in the search display were independently manipulated. For example, among all trials in which the cued object in memory was on the left at encoding (left in internal memory space), this object would equally often occur on the left, right, top, or bottom in the search display. This critical aspect of our design ensured that the relevant internal and external locations during search were fully independent across trials, enabling us to independently track internal and external selection through spatial modulations in gaze and neural activity.

As shown in **Supplementary Figure 1**, Experiment 1 used images of real-world objects (hat, glass, tennis ball, chess piece; around 2 × 2 visual degrees) in the colors: green (RGB: 124, 155, 77), blue (RGB: 97, 154, 152), purple (RGB: 144, 111, 195), and red (RGB: 177, 113, 113). These objects were selected from a public image pool designed for vision experiments (https://bradylab.ucsd.edu/stimuli.html, see also ^53^). Experiment 2 used simple geometric shapes (circle, square, triangle, diamond; matched to have a surface size of 2.25 square visual degrees) in the colors green (RGB: 133, 194, 18), purple (RGB: 197, 21, 234), orange (RGB: 234, 74, 21), and blue (RGB: 21, 165, 234). The choice of specific objects, their colours, and their positions were counterbalanced across trials. The only constraint was that during memory encoding, the two memory objects always occupied opposite locations (left and right or top and bottom). The cued memory object appeared equally often in all four directions in the encoding display and in all four locations of the search display (with its location being independent between encoding and search).

In Experiment 1, participants completed two sessions of 8 blocks, each containing 48 trials, resulting in 768 trials (192 trials per direction in both the internal and the external layout) per participant. The first part of Experiment 2 mirrored the number of trials in Experiment 1. Because the second part of experiment 2 added additional EEG measurements, we increased the number of trials while leaving the experiment unchanged: participants completed three sessions of 8 blocks, each containing 64 trials, resulting in 1536 trials (384 trials per direction in both the internal and the external layout) per participant.

The experiments were programmed in Python (version 3.6.13) with psychopy (version 2021.2.2). During the experiments, participants sat in front of a monitor at a viewing distance of approximately 70 cm with their heads resting on a chin rest.

### Analysis of behavioural performance

Task accuracy and response times were analysed to evaluate performance. Trials in which responses occurred after the search display disappeared were excluded from the analysis. Response times were trimmed to exclude outliers using 2.5 standard deviations beyond the participant’s mean as the cut-off. In both experiments, 99% of the trials were preserved following this trimming.

### Eye-tracking acquisition and pre-processing

A single eye was recorded at 1000 Hz using an EyeLink 1000 system (SR Research). The eye-tracker camera was positioned approximately 5 cm in front of the monitor, 65 cm away from the participant. Built-in calibration and validation procedures were employed before starting the task. Eye-tracking data were analysed offline, in MATLAB using custom scripts (building on ^29^) and the FieldTrip toolbox ^54^. Before turning to saccade detection, we identified blinks by detecting 0 clusters in the gaze data, and then we set all data from 100 ms before to 100 ms after the detected 0 clusters to Not-a-Number (NaN) to remove any residual blink artifacts. Data were then epoched from 1000 ms before to 1500 ms after cue onset.

### Gaze shift detection

Saccades were identified using a velocity-based detection method that we previously validated (^29,55^) and extensively used in complementary prior studies employing the same gaze marker to track internal selection within the spatial layout of visual working memory ^28–31,56–58^.

Gaze velocity was calculated as the Euclidean distance between temporally successive gaze-position values in the 2-dimensional plane (horizontal and vertical gaze position). To improve signal-to-noise ratio (SNR), velocity was smoothed with a Gaussian-weighted moving average filter (7-ms sliding window) using MATLAB’s *smoothdata* function. Saccades were detected when velocity exceeded a trial-specific threshold of 5 times the median velocity, with the first sample crossing the threshold marked as saccade onset. To avoid double counting of saccades, a minimum interval of 100 ms was enforced between successive saccade classifications. The magnitude and direction of each saccade were determined by comparing pre-saccade gaze positions (−50 to 0 ms relative to saccade onset) with post-saccade gaze positions (50 to 100 ms after saccade onset).

We focused our analysis on saccades that moved gaze away from central fixation, defined as saccades whose post-saccade distance to central fixation exceeded the pre-saccade distance. To deal with potential drift after initial calibration, gaze positions corresponding to central fixation were defined empirically at the level of individual trials, as the median gaze position during the fixation period spanning −0.8 to −0.2 ms relative to cue onset.

Gaze shift rates (in Hz) were computed using a 100 ms sliding window advanced in 1-ms steps. To quantify spatial saccade biases, saccades were classified as *toward* or *away* from the location of cued memory object as defined by its location at memory encoding (internal selection) and its location in the search display (external selection). Because the encoded location of the cued memory object was independently varied from its location in the search display, we could independently define spatial biases associated with internal and external selection.

### EEG acquisition and pre-processing

EEG data were recorded at 1024 Hz using a 64-channel BioSemi ActiveTwo system with a conventional 10–10 electrode setup together with left and right mastoids for offline re-referencing. Electro-oculogram (EOG) electrodes were placed near the eyes (horizontally at the left and right eyes, and vertically above and below the left eye) to identify blink and saccade artifacts during Independent Component Analysis (ICA).

EEG analysis was performed in MATLAB (2024b) using the FieldTrip toolbox ^54^ and custom scripts. Data were epoched from 1000 ms before to 1500 ms after cue onset and re-referenced to the mastoid average. ICA was applied, and components correlating with EOG signals were identified and removed. Trials with exceptionally high variance were subsequently rejected using *ft_rejectvisual* in FieldTrip. All data cleaning steps were conducted blind to the experimental conditions of interest. After cleaning, we had 1369 trials (89% of all trials) left for our EEG analysis.

### Neural decoding of internal and external selection

Multivariate decoding was conducted on raw EEG responses that we baseline-corrected using a 250-ms pre-cue baseline window. To decode internal and external selection, we decoded, respectively, the direction of the cued memory object relative to fixation in the encoding display (internal selection) or its direction in the external search display (external selection). Because locations were independently drawn, these decoders were uncorrelated by virtue of our experimental design.

Decoding at each time point used the multivariate Mahalanobis distance (as in ^35,38,59,60^), with electrodes as dimensions, using a leave-one-out approach. For each trial, distances to the average patterns of matching and non-matching classes (defined by the internal/external location of the cued target) were calculated. If multivariate neural patterns contained class-relevant information, distances to the matching class should be smaller than to the non-matching class. Decoding scores were expressed as positive values by subtracting the multivariate Mahalanobis distance with the left-out non-matching trials from the distance with the left-out matching trials. We employed two-class classifiers who were always trained to distinguish target locations in opposite directions (left vs. right or top vs. right) within the memory and search displays. To facilitate visualization, trial-averaged decoding time courses were lightly smoothed with a 100 ms boxcar window (following the same approach we adopted when calculating the rate of gaze shift).

In addition to the above analysis where we always trained and tested the classifier within either internal or external domains, we also ran a cross-domain decoding analysis where we trained the classifier on the direction in one domain and tested it on either the same domain (train-internal->test-internal, train-external->test-external) or the other domain (train-internal->test-external; train-external->test-internal). This enabled us to assess to what extent the neural activity patterns reflecting internal and external selection were shared vs. distinct.

To zoom in on visual activity, the primary decoding analysis utilized a-priori defined cluster of posterior electrodes (O1, PO3, PO7, P1, P3, P5, P7, O2, PO4, PO8, P2, P4, P6, P8) as in ^29^. Additionally, decoding topographies were reconstructed using a searchlight approach ^11,35,38^, iteratively running the same decoding pipeline for subsets of electrodes. For this, we first applied a Laplacian-transform ^61^ to increase spatial resolution and down-sampled the data to 100Hz to increase the decoding efficiency for this more demanding search-light computation. The searchlight consisted was always centred on one electrode and incorporated its immediate lateral neighbours (as in ^11,35,38^).

### Statistical analysis

For all time-series data, we applied a cluster-based permutation approach ^62^ to evaluate the reliability of neural and gaze patterns across neighbouring time points while controlling for multiple comparisons. A permutation distribution of the largest cluster that would occur by chance was generated by randomly permuting the trial-averaged, condition-specific data at the participant level. For each cluster observed in the original data, p-values were calculated as the proportion of permutations where the largest cluster exceeded the size of the observed cluster in the non-permuted data. We performed 10,000 permutations using Fieldtrip’s default settings: clusters were formed by grouping adjacent same-signed data points significant in a mass univariate *t*-test (two-sided alpha = 0.05), with cluster size defined as the sum of *t*-values within the cluster.

To compare the latency of spatial saccade biases between internal and external selection, we additionally used a jackknife approach ^37^. Onset latency was defined as the time point when the saccade bias first reached 50% of its peak value. Latency differences were assessed using the jackknife method described in ^37^ and statistically evaluated using both paired-samples *t*-tests and Bayesian analyses serving to quantify evidence in favour of the null hypothesis (BF01).

Behavioral performance measures (accuracy and reaction time) were also statistically compared using paired-sample *t*-tests supplemented with Bayesian analyses, with reported Bayes Factors expressed as evidence in favour of the null hypothesis of no difference (BF01).

## ACKNOWLEDGEMENTS

This work was supported by an ERC Starting Grant from the European Research Council (MEMTICIPATION, 850636) and an NWO Vidi Grant from the Dutch Research Council (14721) to F.v.E. The authors also thank Chris Olivers, Daniela Gresch, and Kia Nobre for valuable discussions and input.

## Competing interests statement

The authors declare no competing interests.

## Data availability

All data will be made publicly available before publication.

## Code availability

Relevant code associated with the analyses presented here will be made available through GitHub before publication.

## SUPPLEMENTARY INFORMATION

**Supplementary Figure 1.**
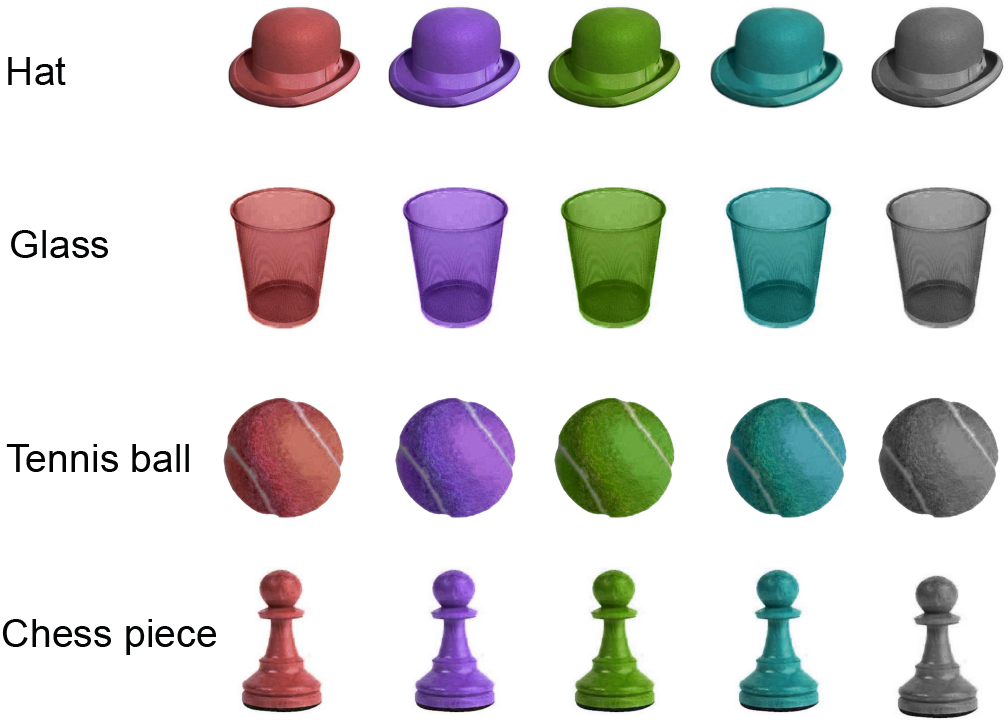
Stimuli used in Experiment 1.

### Control analyses

Here we rule out two possible low-level explanations for the observed concurrence of internal and external selection in our task. Because this involved follow-up control analyses on subsets of our trials, we focused these controls on Experiment 2 where we had more trials and participants available.

First, even though internal and external locations were independently manipulated in our task, it is conceivable that the reported concurrence is exclusively driven by those trials where the internal and external selection targets shared the same direction. To rule this out, we performed a follow-up analysis (control 1) where we exclusively considered trials in which the cued memory object and the matching search target were in different axes at perpendicular directions (e.g. memory object right, matching search target top). We still observed the same concurrent unfolding of our internal and external selection signals (**Supplementary Fig. 2a** jackknife analysis: p = 0.81, BF01 = 6.3).

Second, even though our tasks required selectively searching for the cued memory objects – not just any memory object – it is conceivable that our external-selection marker partly reflects attention being drawn to anything in the search display that looks familiar (i.e., to both memory objects). To rule out any influence of such familiarity, we performed another follow-up analysis (control 2). This time, we exclusively considered trials where the two memory objects *competed* along the same axis in the search display. This ensured that our external-selection signal – reflecting the difference in saccades toward vs. away from the *cued* search target – was exclusively sensitive to selection of the cued memory object (not just any memory object) on the screen. As with control 1, we still observed the same concurrent unfolding of our internal and external selection signals (**Supplementary Fig. 2b**; jackknife analysis: p = 0.51, BF01 = 5.3).

**Supplementary Figure 2.**
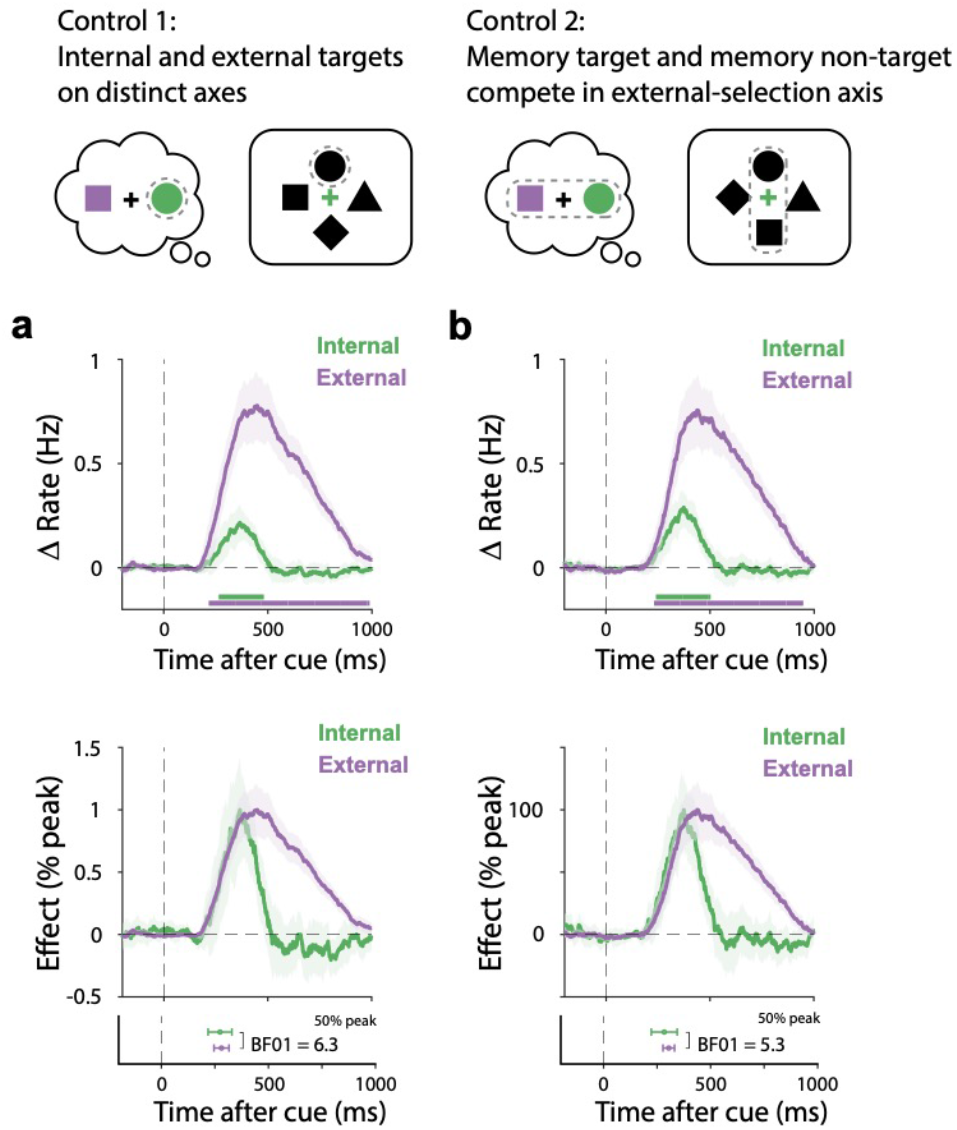
Control analyses rule out low-level explanations for concurrent selection. **a)** Time courses of spatial saccade modulations associated with internal and external selection in trials with perpendicular internal and external target directions. **b)** Time courses of spatial saccade modulations associated with internal and external selection in trials where the two memory objects competed along the same axis in the search display. Bottom panels show peak-normalised data together with onset-latencies calculated as the first sample reaching 50% of the peak. All time courses show mean values, with shading indicating ±1 SEM calculated across participants. The thick horizontal lines in the time course plots indicate significant temporal clusters (cluster-based permutation P < 0.05 (ref)). Error bars on the onset latencies in the bottom panels were estimated using a Jackknife approach and show mean ± the 95% confidence interval. Bayes factors indicate evidence in favour of the null hypothesis of no difference.

## Notes

### Competing Interest Statement

The authors have declared no competing interest.

## REFERENCES

1. Awh, E., Belopolsky, A. V. & Theeuwes, J. Top-down versus bottom-up attentional control: a failed theoretical dichotomy. Trends in Cognitive Sciences 16, 437–443 (2012).

2. Carrasco, M. Visual attention: The past 25 years. Vision Res. 51, 1484–1525 (2011).

3. Desimone, R. & Duncan, J. Neural mechanisms of selective visual attention. Annu. Rev. Neurosci. 18, 193–222 (1995).

4. Moore, T. & Zirnsak, M. Neural Mechanisms of Selective Visual Attention. Annual Review of Psychology vol. 68 47–72 (2017).

5. Wolfe, J. M. Visual Search: How Do We Find What We Are Looking For? Annual Review of Vision Science vol. 6 539–562 (2020).

6. Griffin, I. C. & Nobre, A. C. Orienting Attention to Locations in Internal Representations. J. Cogn. Neurosci. 15, 1176–1194 (2003).

7. Panichello, M. F. & Buschman, T. J. Shared mechanisms underlie the control of working memory and attention. Nature 592, 601–605 (2021).

8. Souza, A. S. & Oberauer, K. In search of the focus of attention in working memory: 13 years of the retro-cue effect. Atten. Percept. Psychophys 78, 1839–1860 (2016).

9. van Ede, F. & Nobre, A. C. Turning Attention Inside Out: How Working Memory Serves Behavior. Annu. Rev. Psychol. (2023) doi:10.1146/annurev-psych-021422-041757.

10. Cavanah, P. J. & Fiebelkorn, I. C. A domain-general process for theta-rhythmic sampling of either environmental information or internally stored information. bioRxiv 2024.11.26.625454 (2024) doi:10.1101/2024.11.26.625454.

11. Gresch, D., Boettcher, S. E. P., Gohil, C., van Ede, F. & Nobre, A. C. Neural dynamics of shifting attention between perception and working-memory contents. Proceedings of the National Academy of Sciences 121, e2406061121 (2024).

12. Verschooren, S., Schindler, S., De Raedt, R. & Pourtois, G. Switching attention from internal to external information processing: A review of the literature and empirical support of the resource sharing account. Psychonomic Bulletin & Review 26, 468–490 (2019).

13. Chun, M. M., Golomb, J. D. & Turk-Browne, N. B. A Taxonomy of External and Internal Attention. Annual Review of Psychology vol. 62 73–101 (2011).

14. Kiyonaga, A. & Egner, T. Working memory as internal attention: Toward an integrative account of internal and external selection processes. Psychonomic Bulletin & Review 20, 228–242 (2013).

15. Smallwood, J. et al. The default mode network in cognition: a topographical perspective. Nature Reviews Neuroscience 22, 503–513 (2021).

16. Spreng, R. N., Stevens, W. D., Chamberlain, J. P., Gilmore, A. W. & Schacter, D. L. Default network activity, coupled with the frontoparietal control network, supports goal-directed cognition. NeuroImage 53, 303–317 (2010).

17. Weilnhammer, V., Stuke, H., Standvoss, K. & Sterzer, P. Sensory processing in humans and mice fluctuates between external and internal modes. PLOS Biology 21, e3002410 (2023).

18. Fiebelkorn, I. C. & Kastner, S. A Rhythmic Theory of Attention. Trends in Cognitive Sciences 23, 87–101 (2019).

19. Honey, C. J., Newman, E. L. & Schapiro, A. C. Switching between internal and external modes: A multiscale learning principle. Network Neuroscience 1, 339–356 (2017).

20. Li, Y. P., Wang, Y., Turk-Browne, N. B., Kuhl, B. A. & Hutchinson, J. B. Perception and memory retrieval states are reflected in distributed patterns of background functional connectivity. NeuroImage 276, 120221 (2023).

21. Poskanzer, C. & Aly, M. Switching between External and Internal Attention in Hippocampal Networks. J. Neurosci. 43, 6538 (2023).

22. Sridharan, D., Levitin, D. J. & Menon, V. A critical role for the right fronto-insular cortex in switching between central-executive and default-mode networks. Proceedings of the National Academy of Sciences 105, 12569–12574 (2008).

23. Treder, M. S. et al. The hippocampus as the switchboard between perception and memory. Proceedings of the National Academy of Sciences 118, e2114171118 (2021).

24. Carlisle, N. B., Arita, J. T., Pardo, D. & Woodman, G. F. Attentional Templates in Visual Working Memory. J. Neurosci. 31, 9315 (2011).

25. Olivers, C. N. L., Peters, J., Houtkamp, R. & Roelfsema, P. R. Different states in visual working memory: when it guides attention and when it does not. Trends Cogn. Sci. 15, 327–334 (2011).

26. Wolfe, J. M. Saved by a Log: How Do Humans Perform Hybrid Visual and Memory Search? Psychol Sci 23, 698–703 (2012).

27. Yu, X., Zhou, Z., Becker, S. I., Boettcher, S. E. P. & Geng, J. J. Good-enough attentional guidance. Trends in Cognitive Sciences 27, 391–403 (2023).

28. Liu, B., Alexopoulou, Z.-S. & van Ede, F. Jointly looking to the past and the future in visual working memory. eLife 12, RP90874 (2024).

29. Liu, B., Nobre, A. C. & van Ede, F. Functional but not obligatory link between microsaccades and neural modulation by covert spatial attention. Nat. Commun. 13, 3503 (2022).

30. van Ede, F., Chekroud, S. R. & Nobre, A. C. Human gaze tracks attentional focusing in memorized visual space. Nat. Hum. Behav. 3, 462–470 (2019).

31. Wang, S. & van Ede, F. Re-focusing visual working memory during expected and unexpected memory tests. eLife 13, RP100532 (2024).

32. Kuo, B.-C., Rao, A., Lepsien, J. & Nobre, A. C. Searching for Targets within the Spatial Layout of Visual Short-Term Memory. J. Neurosci. 29, 8032 (2009).

33. Luck, S. J. & Hillyard, S. A. Electrophysiological correlates of feature analysis during visual search. Psychophysiology 31, 291–308 (1994).

34. Schneider, D., Barth, A. & Wascher, E. On the contribution of motor planning to the retroactive cuing benefit in working memory: Evidence by mu and beta oscillatory activity in the EEG. Neuroimage 162, 73–85 (2017).

35. van Ede, F., Chekroud, S. R., Stokes, M. G. & Nobre, A. C. Concurrent visual and motor selection during visual working memory guided action. Nat. Neurosci. 22, 477–483 (2019).

36. van Moorselaar, D., Gunseli, E., Theeuwes, J. & N. L. Olivers, C. The time course of protecting a visual memory representation from perceptual interference. Frontiers in Human Neuroscience 8, (2015).

37. Smulders, F. T. Y. Simplifying jackknifing of ERPs and getting more out of it: Retrieving estimates of participants’ latencies. Psychophysiology 47, 387–392 (2010).

38. Kandemir, G. & Olivers, C. Comparing Neural Correlates of Memory Encoding and Maintenance for Foveal and Peripheral Stimuli. Journal of Cognitive Neuroscience 36, 1807–1826 (2024).

39. Kriegeskorte, N., Goebel, R. & Bandettini, P. Information-based functional brain mapping. Proceedings of the National Academy of Sciences 103, 3863–3868 (2006).

40. Zhou, Y. (周颖), Curtis, C. E., Sreenivasan, K. K. & Fougnie, D. Common Neural Mechanisms Control Attention and Working Memory. J. Neurosci. 42, 7110 (2022).

41. Weber, R. J., Burt, D. B. & Noll, N. C. Attention switching between perception and memory. Memory & Cognition 14, 238–245 (1986).

42. Soto, D., Hodsoll, J., Rotshtein, P. & Humphreys, G. W. Automatic guidance of attention from working memory. Trends in Cognitive Sciences 12, 342–348 (2008).

43. Folk, C. L., Remington, R. W. & Johnston, J. C. Involuntary covert orienting is contingent on attentional control settings. Journal of Experimental Psychology: Human perception and performance 18, 1030 (1992).

44. Downing, P. E. Interactions Between Visual Working Memory and Selective Attention. Psychol Sci 11, 467–473 (2000).

45. Awh, E. & Jonides, J. Overlapping mechanisms of attention and spatial working memory. Trends in Cognitive Sciences 5, 119–126 (2001).

46. Serences, J. T. et al. Coordination of Voluntary and Stimulus-Driven Attentional Control in Human Cortex. Psychol Sci 16, 114–122 (2005).

47. Dell’Acqua, R., Sessa, P., Toffanin, P., Luria, R. & Jolicœur, P. Orienting attention to objects in visual short-term memory. Neuropsychologia 48, 419–428 (2010).

48. Liu, B., Kong, S. & van Ede, F. Microsaccades strongly modulate but do not cause the N2pc EEG marker of spatial attention. bioRxiv 2024.10.28.620656 (2024) doi:10.1101/2024.10.28.620656.

49. Verschooren, S., Schindler, S., De Raedt, R. & Pourtois, G. Early reduction of sensory processing within the visual cortex when switching from internal to external attention. Biological Psychology 163, 108119 (2021).

50. Libby, A. & Buschman, T. J. Rotational dynamics reduce interference between sensory and memory representations. Nature Neuroscience 24, 715–726 (2021).

51. Xu, Y. The human posterior parietal cortices orthogonalize the representation of different streams of information concurrently coded in visual working memory. PLOS Biology 22, e3002915 (2024).

52. Myers, N. E. et al. Testing sensory evidence against mnemonic templates. eLife 4, e09000 (2015).

53. Brady, T. F., Konkle, T., Gill, J., Oliva, A. & Alvarez, G. A. Visual Long-Term Memory Has the Same Limit on Fidelity as Visual Working Memory. Psychol Sci 24, 981–990 (2013).

54. Oostenveld, R., Fries, P., Maris, E. & Schoffelen, J.-M. FieldTrip: Open Source Software for Advanced Analysis of MEG, EEG, and Invasive Electrophysiological Data. Comput. Intel. Neurosc. 2011, 156869 (2011).

55. Liu, B., Nobre, A. C. & van Ede, F. Microsaccades transiently lateralise EEG alpha activity. Prog. Neurobiol. 224, 102433 (2023).

56. Chawoush, B., Draschkow, D. & van Ede, F. Capacity and selection in immersive visual working memory following naturalistic object disappearance. J. Vis. 23, 9–9 (2023).

57. de Vries, E., Fejer, G. & van Ede, F. No obligatory trade-off between the use of space and time for working memory. Commun. Psychol. 1, 41 (2023).

58. van Ede, F., Board, A. G. & Nobre, A. C. Goal-directed and stimulus-driven selection of internal representations. Proc. Natl. Acad. Sci. U.S.A. 117, 24590–24598 (2020).

59. van Ede, F., Chekroud, S. R., Stokes, M. G. & Nobre, A. C. Decoding the influence of anticipatory states on visual perception in the presence of temporal distractors. Nature Communications 9, 1449 (2018).

60. Wolff, M. J., Jochim, J., Akyürek, E. G. & Stokes, M. G. Dynamic hidden states underlying working-memory-guided behavior. Nature Neuroscience 20, 864–871 (2017).

61. Perrin, F., Pernier, J., Bertrand, O. & Echallier, J. F. Spherical splines for scalp potential and current density mapping. Electroencephalography and Clinical Neurophysiology 72, 184–187 (1989).

62. Maris, E. & Oostenveld, R. Nonparametric statistical testing of EEG- and MEG-data. J. Neurosci. Methods 164, 177–190 (2007).

